# Highly Consistent Anatomical Asymmetry in a Small Primate Brain: Left is Always Larger in the Marmoset Monkey

**DOI:** 10.1101/2025.11.19.689300

**Authors:** Inaki C. Mundinano, Nafiseh Atapour, Kun Jiang, Ranshikha Samandra, Cirong Liu, Farshad Mansouri, Marcello G. P. Rosa

## Abstract

Although lateralisation of brain function is relatively common in animals, this is rarely accompanied by overt anatomical asymmetries. In primates, morphological differences between the cerebral hemispheres are well established in Hominidae but have been regarded as subtler or inconsistent in other species. Here we demonstrate that the left hemisphere is reproducibly larger than the right hemisphere in one of the smallest primates, the marmoset. This asymmetry develops postnatally, persists throughout adult life, and is found in individuals from genetically isolated colonies. Voxel-based morphometry reveals that the larger left hemisphere is primarily due to differences in the volume of cortical areas linked to social cognition. This result challenges the notion that the development and evolution of marked morphological asymmetry between the hemispheres is linked to the evolution of large brains.

---

Although lateralisation of brain function is common across the animal kingdom ^1^, this is not usually linked to morphological differences between the two cerebral hemispheres. For example, although many species show lateralisation according to criteria such as the preferred use of one of the limbs ^2^, development and production of communication calls ^3,4^, spatial cognition ^5^ and motor expression of emotional cues ^6,7^, this is rarely accompanied by overt anatomical asymmetries. Most notably, humans and apes show consistent morphological differences including (but not restricted to) cortical areas related to the evolution of language ^8-10^. But even this trend is less marked and inconsistent across individuals in other non-human primates such as macaques and baboons, despite functional evidence of hemispheric bias at individual level ^11-13^. This, together with the rarity of asymmetry in other mammalian species, has led to the idea that reproducible changes in the gross anatomy of the cortex may be linked to the evolution of larger brains ^14,15^. In a nutshell, the development of a larger brain would impose constraints related to the efficiency of axonal connections and availability of synaptic space in single neurons; this would lead to the gradual segregation of neural structures into more specialised subnetworks, and ultimately morphological asymmetries. In other words, increasing evolutionary pressure for functional specialisation would only in selected instances be expressed as consistent anatomical differences.

We explored this issue using high resolution magnetic resonance imaging (MRI) in marmoset monkeys (*Callithrix jacchus*), which are among the smallest primates, with a brain volume of only 8g ^16^. Marmosets have become increasingly used as an animal model in the neurosciences due to its highly social behaviour ^17^, which includes a rich repertoire of vocal communication ^18^, as well as a well-developed visual system ^19^ and relatively short reproductive cycle, which facilitates the generation of transgenic lines ^20,21^. Like other mammals they are known to show behavioural evidence lateralisation of motor and sensory function, which, however, varies across individuals ^22,23^.

The primary sample included MRI scans obtained from 302 marmosets obtained either for the purpose of ongoing studies in our laboratories or from publicly available databases. The dataset originated from 4 geographically isolated colonies, located in Australia (Monash; n=9), China (ION; n=60), Japan (RIKEN; n=206) and the USA (NIH; n=17). This included both i*n vivo* (Australia, Japan, USA) and *ex vivo* scans (China). We found that adult marmosets (>2 years old, n= 208; Supplementary Table S1) show a remarkably consistent anatomical asymmetry, whereby the cerebral cortex in left hemisphere is invariably larger than that in the right hemisphere (Fig. 1a). When quantified according to a commonly used metric, the hemispheric asymmetry index (AI), the left hemisphere is on average 2.31% larger than the right (range 0.22%-4.37%), corresponding to a difference in volume of 96.89 ± 27.18 mm^3^, (Wilcoxon Signed-Rank Test: W = 0, p = 7.0131e−36; paired t-test: t = 51.42, p = 7.54e−120; Cohen’s d = 3.57). Male and female marmosets did not differ significantly in this respect (Mann–Whitney U test: *U* = 4203.0000, *p* = 7.8553e−02; rank-biserial correlation: *r* = 0.1549) (Fig. 1b). Data from the different colonies demonstrate the same trend, despite the individuals therein being genetically isolated for at least 10 generations. While statistical differences in the AI distribution were observed between colonies (Kruskal– Wallis: H = 52.4961, p = 2.3478e−11; *ε*^2^ = 0.2067), AI was consistently positive, indicating that leftward hemispheric asymmetry is a universal feature in adult marmosets irrespective of sex, age, or colony of origin (Supplementary Fig. S2).

**Figure 1.**
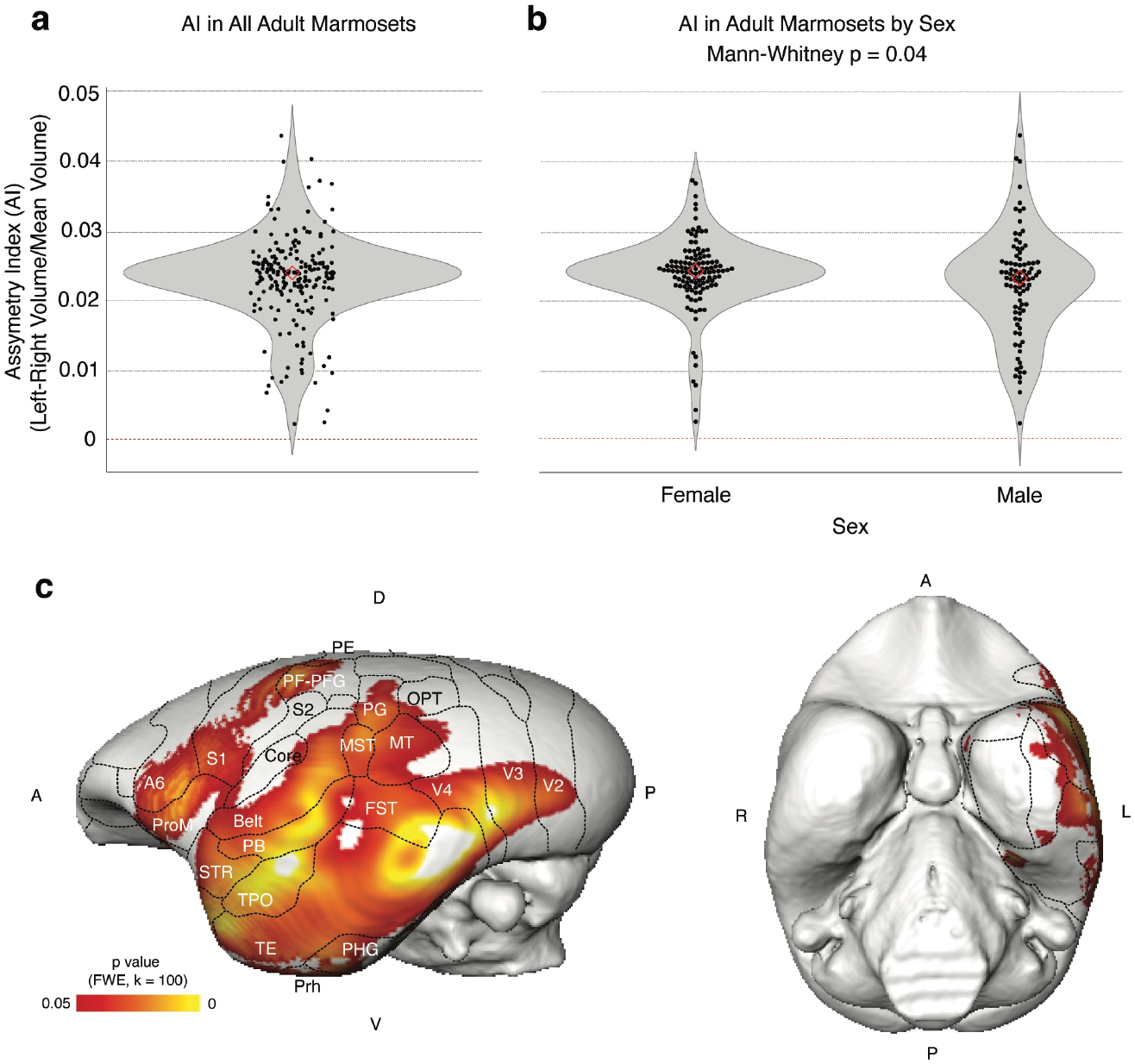
Distribution of hemispheric asymmetry in adult marmosets. **a**: Violin plots of the distribution of the hemispheric Asymmetry Index (AI) in all adult marmosets (>2 years old) included in the study (n = 208). AI quantifies the relative volume between brain hemispheres, with positive values indicating leftward asymmetry. Each dot represents an individual subject, with red diamond outlines indicating group medians. **b**: Distribution of AI values according to sex. Both male (n = 90) and female (n = 118) marmosets exhibit positive AI values, indicating leftward hemispheric asymmetry. **c**: Voxel-based morphometry (VBM) analysis comparing grey matter volume between hemispheres in adult marmosets (n = 208). Statistical maps (yellow-red) indicate cortical regions where grey matter volume is significantly greater in the left hemisphere compared to the right, thresholded at p < 0.05 (FWE-corrected), with a minimum cluster extent of 100 voxels (*k* = 100). The left panel shows a lateral view of the left hemisphere, while the right panel shows a ventral view. Overlaid anatomical boundaries correspond to the MBMv3_M cortical parcellation ^24^. Anatomical orientation: A-anterior, D-dorsal, L-left, P-posterior, R-right, V-ventral. For abbreviations of the names of cortical areas, see ^30^.

The hemispheric asymmetry is not present throughout the brain. Voxel-based morphometry according to registration to a MR-based template ^24^, revealed that the differences are attributed to larger left hemisphere volumes in the visual and polysensory areas of the lateral and inferior temporal cortical regions (cytoarchitectural areas TE, TPO, IPa/ PGa), the high-order auditory cortex (belt, parabelt, and the rostral superior temporal auditory association area), perirhinal cortex, and the foveal representations of some extrastriate visual areas (V3, V4, and MST) (Fig. 1c). Further asymmetries were observed in the frontal lobe (ventral preomotor area, and precentral opercular cortex), and in the ventral posterior parietal cortex (areas PG PFG, PF).

Considering these observations, a natural question is whether the hemispheric asymmetry is present from birth or develops during postnatal life. For this we studied additional individuals (Supplementary Tables S3 and S3), including animals within the first 12 months of postnatal life (n=64), and peri pubescent animals (12-24 months; n=30), and further segregated the adult cohort according to age (Fig. 2a). We found that there is a slight, but inconsistent trend towards a larger left hemisphere already in the youngest animals. The left hemisphere bias becomes consolidated into the emergence of a marked asymmetry during puberty. This is maintained throughout adulthood, including old age (Fig. 2b).

**Figure 2.**
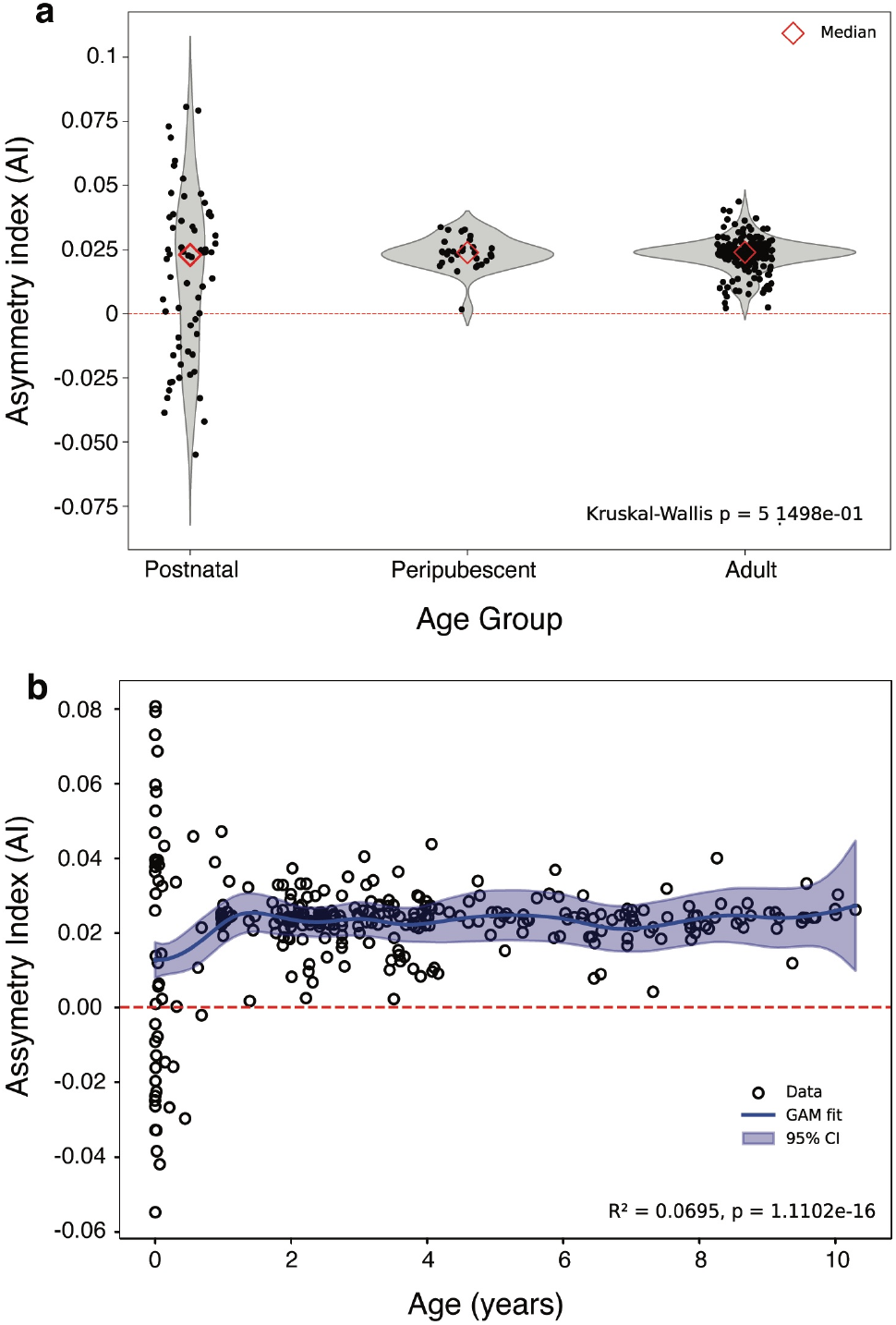
Age-related differences in cerebral asymmetry across the lifespan in marmosets. The asymmetry index (AI) according to developmental stages in marmosets. Postnatal: <1 year old (n = 64). Peripubescent: 1–2 years (n = 30). Adult: >2 years (n = 208). Marmosets are typically considered to be of old age beyond 7-8 years and rarely survive beyond 12 years, even in controlled colony conditions. **a:** All groups demonstrate leftward hemispheric asymmetry (positive AI values), but there is higher variability in the Postnatal group. **b:** Relationship between age and the cortical asymmetry index (AI) across individual marmosets, using a generalized additive model (GAM) to account for potential nonlinear trends. Each point represents one animal. A red dashed horizontal line indicates equal volume in the 2 hemispheres (AI= 0). The model captures subtle age-related patterns in asymmetry, particularly during early development, while highlighting the overall stability of AI across later stages of life.

Perhaps the most surprising aspect of our data is the high reproducibility of the anatomical bias towards the left hemisphere. This finding contrasts with the results reported in most mammals to date, where functional and anatomical asymmetries, when present, tend to be variable across individuals. Although the full extent of leftward hemispheric asymmetry in primates clearly requires more research, what is currently known makes unlikely that the consistently larger left hemispheres observed in both humans and marmosets represent a symplesiomorphic evolutionary trait. Rather, this may represent a case of parallel evolution.

Among the most consistently asymmetric areas in the marmoset are those involved in higher-order visual and auditory recognition and integration of multisensory information, and those involved in olfactory processing and episodic memory. These are all important requirements for social interaction, which are distinctively developed in marmoset groups ^25^. In the frontal and parietal regions, the only areas showing asymmetry are those linked to the “mirror neuron” system in other primate species ^26^. It is therefore possible that the allocation of increased neural resources for social interaction has been an evolutionary driver of functional specialisation, followed by asymmetry in cortical structure and function, in both callitrichids and hominoids. Postnatal developmental mechanisms including hormonal influences and epigenetic modification could account for this, given the strong influence of social context in marmoset behaviour.

Our results reveal that consistent anatomical asymmetries are not an exclusive domain of larger brains, calling into question the association between these and brain volume and the inability of the corpus callosum to maintain proportional connectivity. Among other implications of this finding, it highlights the need to consider laterality whenever anatomical or physiological data obtained in various animal models. Many studies do not report whether observations in non-human primates originated in the left or right hemispheres, or treat observations obtained on both sides of the brain as interchangeable. Moving forward, this is an issue that will require additional attention, possibly including revised editorial guidelines for publication in comparative neuroscience.

## Acknowledgements

We acknowledge the Brain/MINDS project (RIKEN Center for Brain Science, Japan) for providing open access to the marmoset MRI dataset, which greatly contributed to this study. The dataset is available at https://dataportal.brainminds.jp/mri-data-page, (Junichi Hata, Ken Nakae, Daisuke Yoshimaru, Hideyuki Okano. Brain/MINDS Marmoset Brain MRI Dataset NA216 and eNA91: (DataID: 4624) doi:https://doi.org/10.24475/bminds.mri.thj.4624.

We also thank Dr Gang Zheng from the Preclinical Unit of Monash Biomedical Imaging for his invaluable assistance with the acquisition of MRI data from the Monash colony, and Xiaojia Zhu and Haotian Yang for assisting in sample preparation and MRI data collection in the ION.

## Funding

National Health and Medical Research Council Ideas grants (APP2011733, APP2019011), and Investigator grant (APP1194206).

Australian Research Council Discovery Projects (DP210103865, DP210101042, DP250101768)

National Science and Technology Major Program, China (Grant No. 2025ZD0219300 and 2022ZD0205000).

National Natural Science Foundation of China (No. 32427802).

Shanghai Key Laboratory of Child Brain and Development (No. 24dz2260100).

## Supplementary Materials

**Supplementary Table S1.** Demographics: adult animals (>2 years old) included in the study.

| Colony | Subjects | Mean_Age<br>(years) | SD_Age | Males | Females |
| --- | --- | --- | --- | --- | --- |
| ION | 3 | 2.72 | 0.64 | 2 | 1 |
| Monash | 9 | 3.31 | 0.63 | 8 | 1 |
| NIH | 15 | 4.70 | 1.84 | 11 | 4 |
| Riken | 181 | 4.94 | 2.41 | 69 | 112 |
| Total | 208 | 3.92 | 1.38 | 90 | 118 |

**Supplementary Figure S1:**
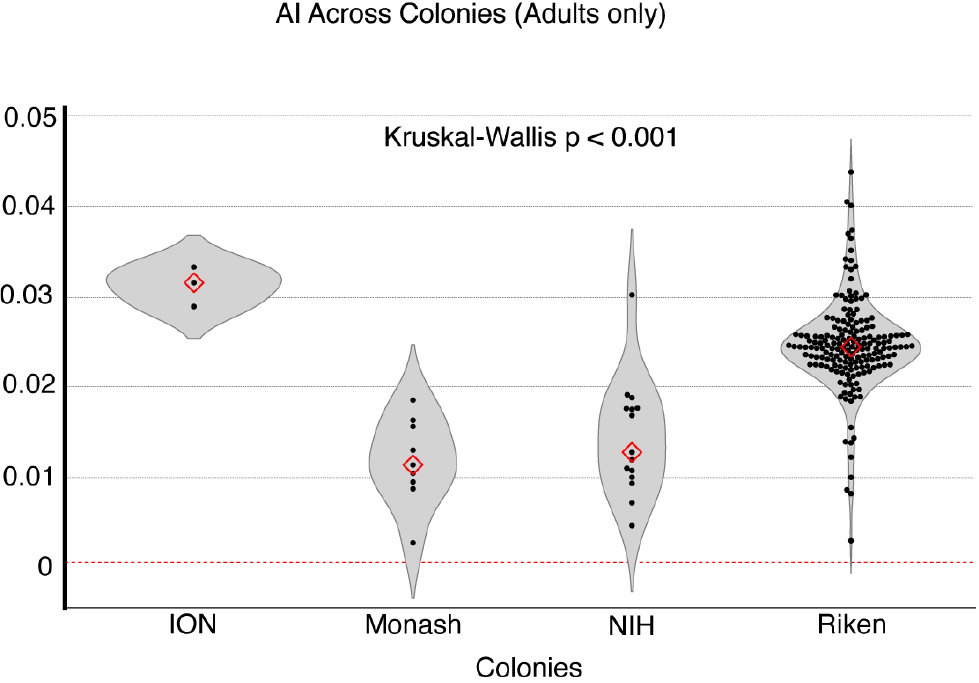
AI values across four independent marmoset colonies in China (ION; n=3), Australia (Monash; n=9), USA (NIH; n=15), and Japan (Riken; n=181). All colonies show consistently positive AI values.

**Supplementary Table S3.** Demographics: Postnatal group (<1 year old)

| Colony | Subjects | Mean_Age<br>(years) | SD_Age | Males | Females |
| --- | --- | --- | --- | --- | --- |
| ION | 53 | 0.14 | 0.24 | 33 | 20 |
| Riken | 11 | 0.98 | 0.01 | 8 | 3 |
| Total | 64 | 0.56 | 0.13 | 41 | 23 |

**Supplementary Table S4.** Demographics:. Peripubescent group (1-2 years old)

| Colony | Subjects | Mean_Age<br>(years) | SD_Age | Males | Females |
| --- | --- | --- | --- | --- | --- |
| ION | 4 | 1.24 | 0.21 | 2 | 2 |
| NIH | 2 | 1.48 | 0.68 | 1 | 1 |
| Riken | 24 | 1.73 | 0.30 | 14 | 10 |
| Total | 30 | 1.48 | 0.40 | 17 | 13 |

## Methods

### Animals and Study Demographics

A total of 302 common marmosets (*Callithrix jacchus*) were included in this study, classified into three age groups: Postnatal (<1 year), Peripubescent (1–2 years), and Adult (>2 years). Supplementary Tables 1, 3 and 4 summarize the demographics by colony, age, and sex. Animals were obtained from four independent colonies: RIKEN Center for Brain Science (Japan), ION (Institute of Neuroscience, China), NIH (National Institutes of Health, USA), and Monash University (Australia). Animals from each colony were bred and group housed according to institutional animal care protocols, each subject to approval by a local Animal Ethics Experimentation Committee.

### MRI acquisition

#### Monash University

MRI data were acquired *in vivo* on a 9.4T Bruker BioSpec scanner equipped with a Bruker 86 mm volume coil for transmission and a Rapid 8-channel array coil for reception. Structural MRI was performed using a two-dimensional T2-weighted Rapid Acquisition with Relaxation Enhancement (RARE) sequence (TR = 3000 ms, TE = 47.4 ms, rare factor = 12, slice thickness = 0.2 mm, FOV = 40 × 40 mm^2^, matrix = 200 × 200, in-plane resolution = 0.2 × 0.2 mm^2^, 100 slices, 3 signal averages, total scan time = 25 min 30 s). Animals were anesthetized with alfaxalone (8 mg/kg, i.m.) and diazepam (3 mg/kg, i.m.), and anaesthesia was maintained with 0.5–2.0% isoflurane during scanning. Animals were positioned in the sphinx posture within a water-heated, MRI-compatible cradle. Body temperature and respiratory rate were continuously monitored throughout the imaging session. During scanning, anaesthesia was maintained with 0.5–2.0% isoflurane delivered in medical-grade oxygen via a custom-fitted face mask.

### ION

*Ex vivo* MRI data for marmoset brain development were acquired at the Chinese Academy of Sciences Institute of Neuroscience (CAS-ION) on a 9.4T Bruker BioSpec 94/30 MRI scanner using a FLASH sequence. Prior to scanning, all brains were preserved *in situ* within the skull and immersed in a solution of 4% paraformaldehyde (PFA) with 0.3% gadolinium (Gd) for 2-4 weeks, followed by a one-week immersion in phosphate-buffered saline (PBS). The dataset comprises two cohorts. For the first cohort (16 male, 10 female) acquisition parameters for this cohort were individually adjusted to account for differences in brain size and to optimize image contrast for animals of varying ages, resulting in a range of values: TR = 20 ms; TE = 3.6 ms to 4.7 ms; matrix size = 225×290×210 to 285×360×270; FOV = 21.6×27.84×20.16 to 22.8×28.8×21.6 mm; flip angle = 15° to 25°; voxel size = 80 μm to 96 μm isotropic; number of average = 1; scan time = 9:57 min to 18:10 min. The second cohort (23 male, 13 female) was scanned with a fixed parameter: TR = 30 ms; TE = 20 ms; matrix = 278×360×270; FOV = 22.2×28.8×21.6 mm; flip angle = 35°; voxel size = 80 μm isotropic; number of average = 2; scan time = 37:58 min.

### RIKEN Center for Brain Science

Images from the from the Japanese colony were obtained from the BRAIN/MINDS marmoset Brain MRI dataset^27^. *In vivo* T2-weighted MRI data were obtained from the BRAIN/MINDS marmoset MRI dataset. Scans were performed using a 9.4T Bruker BioSpec 94/30 MRI scanner (TR = 4000 ms, TE = 22 ms, RARE factor = 4, voxel size = 270 × 270 × 540 ^µ^m, scan time = 7 min 24 s). Animals were anesthetized with 2% isoflurane in an oxygen-air mixture and monitored throughout.

### NIH

*In vivo* MRI data were acquired as described previously ^24^. Briefly, animals were anesthetized with an intramuscular injection of 10 mg/kg ketamine and ventilated with 1.5% - 2% isoflurane. The animals were placed in an MR-compatible cradle, and their vital signs were monitored throughout the MRI scanning. MRI scanning was performed in a Bruker 7T/300 mm magnet with a 2D RARE sequence: TR = 30 s, TE = 8 s with three effective TE (TE1 = 16 ms, TE2 = 48 ms and TE3 = 80 ms), FOV = 36 × 28 × 24 mm, matrix size = 144 × 112,slice thickness = 0.25 mm, and the scanning duration was about 10 min 30 s.

### Image Preprocessing and Volume calculation

All MRI data were processed using a standardized pipeline implemented in Advanced Normalization Tools (ANTs) and the FMRIB Software Library (FSL) ^28^. T2-Weighted images were first skull stripped using the template-based brain extraction tool *atlasBREX* ^29^, using the Marmoset Brain Mapping V3 (MBMv3) atlas ^24^ as a reference to the areas defined by Paxinos et al ^30^. Brain-extracted images were denoised (*DenoiseImage*, Rician noise model, spatial radius = 3), followed by N4 bias field correction (*N4BiasFieldCorrection*, shrink factor = 3, B-spline fitting). Importantly, in order to preserve natural hemispheric brain asymmetry, images were registered to the MBMv3 marmoset atlas space using only affine rigid-body alignment (*antsRegistrationSyNQuick*.*sh*, transform type: rigid) avoiding any non-linear deformation. Binary brain masks were generated using fslmaths, and hemisphere-specific volumes were computed using fslstats with predefined left and right hemisphere masks from the MBMv3 atlas.

### Asymmetry Index Calculation

Hemispheric asymmetry was quantified using the Asymmetry Index (AI):

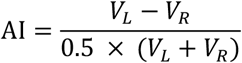

where *V*_*L*_ and *V*_*R*_ represent the left and right hemisphere volumes, respectively. This metric normalizes inter-hemispheric differences by the mean total hemispheric volume. Positive AI values indicate leftward asymmetry (larger left hemisphere), while negative values indicate rightward asymmetry.

### Voxel Based Morphometry

To assess which specific cortical areas contributed to the hemispheric asymmetries in gray matter (GM) volume, a paired voxel-based morphometry (VBM) analysis was employed using SPM12 (Wellcome Trust Centre for Neuroimaging, London, UK) in MATLAB. Preprocessed images were resampled to 1×1×1 mm^3^ isotropic resolution. Then, images were left–right flipped along the x-axis using fslswapdim to generate mirrored versions of each brain. Flipped images were then registered again to their original unflipped images to ensure anatomical correspondence and segmented into grey matter, white matter and cerebrospinal fluid in SPM12 (old segmentation). Gray matter maps were then normalized to the MBMv3 marmoset template using DARTEL, with Jacobian modulation applied to preserve local volumetric information. Smoothed GM maps (6×6×6 mm FWHM) were compared using a voxel-wise paired t-test between original and flipped images. Results were thresholded at p < 0.05, family-wise error (FWE) corrected, with a cluster extent threshold of 100 voxels. Significant values were displayed on the MBMv3 T2-template and cortical areas identified with the MBM_cortex_vM atlas labels.

### Statistical methods

All statistical analyses were conducted using Python (version 3.10.15) *scipy, statsmodels, scikit-posthocs and pyGAM* libraries. To assess whether the left hemisphere was significantly larger than the right in adult marmosets, we used the Wilcoxon Signed-Rank Test, a non-parametric paired test. Differences in the asymmetry index (AI) between colonies and age groups were analysed using the Kruskal–Wallis test, followed by Dunn’s post-hoc test with Bonferroni correction to evaluate pairwise group differences. To examine sex differences in AI, we employed the Mann–Whitney U test. Additionally, we investigated the relationship between age and AI using both linear regression and a non-parametric Generalized Additive Model (GAM). The GAM framework allowed for the detection of potential nonlinear age-related trends in asymmetry by fitting a smooth spline function to the data. To evaluate the combined effects of age, sex, and colony on inter-hemispheric volume differences, we employed multiple linear regression using the *statsmodels*.*OLS* framework. Effect sizes were calculated to complement statistical significance testing, including epsilon-squared (ε^2^) for Kruskal–Wallis comparisons and rank-biserial correlation for the Mann–Whitney U test. All results were considered statistically significant at p < 0.05. Visualizations were generated using the *matplotlib* and *seaborn* libraries, employing violin plots overlaid with individual data points (swarm/strip plots) and red diamond markers to indicate median values.

